# miREA: a network-based tool for microRNA-oriented enrichment analysis

**DOI:** 10.64898/2026.02.27.708509

**Authors:** Zhesi Zhang, Xin Lai

**Author notes:** Corresponding author: Xin Lai, Faculty of Medicine and Health Technology, Tampere University, Tampere, Finland. **Data and code availability** Zenodo: https://doi.org/10.5281/zenodo.18803209, GitHub: https://github.com/laixn/miREA.

## Abstract

MicroRNAs (miRNAs) regulate gene expression at the post-transcriptional level, yet interpreting their function at the pathway level remains challenging. Existing enrichment analysis tools predominantly adopt network node-centric approaches that focus on gene expression profiles, neglecting the regulatory information encoded in miRNA-gene interactions (MGIs) that constitute network edges. This omission introduces analytical bias and limits biological interpretability, underscoring the need for network edge-based enrichment analysis methods that explicitly incorporate MGIs. Therefore, we present miREA, a network-based tool for miRNA enrichment analysis that leverages MGIs to characterize miRNA function at the pathway level. miREA contains five edge-based enrichment methods that integrate paired miRNA-gene expression and interactome profiles with pathway networks to perform MGI overrepresentation, MGI scoring-based, network topology-aware, and network propagation analyses. Benchmarking across multiple cancer types shows that the edge-based methods outperform node-based methods in improving sensitivity to identify relevant pathways and biological interpretability while maintaining controlled false positive rates. We further demonstrate the utility of miREA in elucidating miRNA-gene-pathway regulatory mechanisms in bladder cancer. miREA is a versatile enrichment analysis tool that provides pathway-level interpretation of human miRNA function and facilitates mechanistic hypothesis generation for experimental validation.

**Highlights:**

- miREA uses information in miRNA-gene interactions for miRNA enrichment analysis.
- miREA contains five new edge-based algorithms that integrate expression and interactome profiles with pathway networks to characterize miRNA function.
- The edge-based methods are more sensitive than node-based methods at identifying cancer-relevant pathways and more effective at identifying cancer genes and miRNAs.
- We perform systematic analysis of the miREA methods to show their robustness on 16 cancer datasets.
- We present a case study that demonstrates the potential of miREA to elucidate miRNA-gene-pathway regulatory mechanisms in bladder cancer.

## 1 INTRODUCTION

MicroRNAs (miRNAs) are a class of small non-coding RNAs that have been shown to participate in diverse biological processes and disease progression (1–3). Specifically, they regulate gene expression at the post-transcriptional level by binding to complementary sequences in target mRNAs, predominantly functioning through translational repression or mRNA degradation (4–6). Unveiling the functional roles of miRNAs is essential for understanding their regulatory mechanisms and discovering potential therapeutic targets. A single miRNA can regulate numerous transcripts involved in diverse cellular functions (7, 8), whereas individual mRNAs may also be targeted by multiple miRNAs (9, 10). This intricate many-to-many regulatory relationship complicates the interpretation of differential miRNA expression and makes it challenging for the functional annotations of individual miRNAs in cellular processes (11–13). Moreover, the development of high-throughput sequencing techniques has produced massive amounts of data, making computational approaches indispensable for the systematic analysis of large-scale miRNA datasets (14, 15). So, developing a versatile tool for analyzing miRNA sequencing data is appealing for the research community.

Enrichment analysis has been widely applied to summarize gene-level or miRNA-level expression profiles to pathway-level biological insights (16–18). A pathway represents a functional molecular set that collectively regulates specific biological processes or cellular functions. Depending on the data utilized and the underlying algorithms, gene set enrichment analysis can be broadly classified into four major categories, including over-representation analysis (ORA) (19), scoring-based analysis (20), pathway topology analysis (21), and network-based analysis (22). Conventional miRNA enrichment analysis are either a gene-centric (23–25) or a miRNA-centric method (16, 26). Gene-centric methods identify genes targeted by miRNAs of interest (e.g., differentially expressed miRNAs of a biological condition) and perform hypergeometric tests on these miRNA target genes to identify their enrichment in relevant pathways. In contrast, miRNA-centric methods infer miRNA-associated pathways either from curated miRNA sets associated with certain diseases and functions (26), or by transforming gene-associated pathways into miRNA sets that target at least one gene within the pathways. These methods can be collectively regarded as node-based approaches, the most representative of which include miEAA (16) and miRNet (17) that provide both gene-centric and miRNA-centric analysis through interactive web interfaces. However, node-based approaches introduce biases by considering the associations between miRNAs and their targets as binary and neglecting the regulatory characteristics of miRNA-gene interactions (MGIs), such as ignorance of miRNA’s binding attributes to their target genes and the regulatory strengths between them (12, 13, 27), thereby downplaying the topological attributes of gene regulatory networks in pathways (28, 29). In contrast, edge-based enrichment analysis methods address these limitations by considering the regulatory information in MGIs, thereby endowing the analysis with more accurate regulatory and topological information in gene regulatory networks. Specifically, differential co-expression analysis has been employed to identify dysregulated pathway gene sets. Han et al. investigates dysregulated pathways using gene expression data through the edge set enrichment analysis that are based on differential correlation analysis of gene-gene interactions (GGIs) (30). DysPIA performs paired gene enrichment analysis by quantifying dysregulated gene pairs based on two-sample Welch’s t-test (31). However, these methods focus exclusively on protein-coding genes and rely heavily on expression profiles than regulation information (32). This limits their abilities to capture regulatory insights and makes them sensitive to variability in transcriptomic data. Moreover, expression-based edge analysis has limited mechanistic interpretability, which is crucial for understanding miRNAs’ role because their function is mainly determined by their interacting target genes (5, 33). These limitations highlight the need for an edge-based enrichment analysis approach that integrates reliable MGI landscape, quantitative interaction information, and curated pathway networks to systematically analyze the function of miRNAs.

We develop a network-based tool for <u>miR</u>NA-oriented <u>E</u>nrichment <u>A</u>nalysis (miREA) that measures miRNAs at the edge level to characterize miRNA regulatory functions related to biological pathways. By focusing on MGIs instead of individual miRNAs or genes, miREA addresses the intrinsic many-to-many MGI regulatory roles and reduces biases introduced by node-centric enrichment approaches, where MGI scores are proposed to measure and differentiate the regulatory strengths of MGIs. miREA integrates paired miRNA and gene expression profiles with curated gene regulatory networks, including MGI and GGI. It implements five edge-based algorithms that perform MGI overrepresentation (Edge-ORA), MGI scoring-based (Edge-Score and Edge-2Ddist), network topology-aware (Edge-Topology), and network propagation (Edge-Network) analyses. Through comparative analysis of node-based methods on 17 cancer types, we show the superiority of the edge-based algorithms in characterizing miRNAs’ functions. These improvements are demonstrated by better metric values, including the sensitivity and specificity of biologically relevant pathways, the ability to distinguish between cancer-specific pathways, and the ability to identify cancer-related miRNAs and genes. Systematic analyses demonstrate the robustness of the methods to input data variance as well as their computational efficiency using parallel computing. In addition, miREA provides advanced visualization to facilitate method selection and investigation of miRNA-gene-pathway regulatory mechanisms, as demonstrated through a bladder cancer case study. Collectively, miREA is a versatile tool for analyzing miRNA function at the pathway level, offering mechanistic insights into miRNA-mediated gene regulation.

## 2 MATERIALS AND METHODS

The complete computational workflow of this study is described in the following sections and illustrated in Figure S1. In brief, the workflow contains several parts, including processing of MGI and GGI, pathway networks, and gene expression data (Sections 2.1-2.3), method development (Sections 2.4 and 2.5), method evaluation (Sections 2.6, 2.7, and 3.2), method summary (Section 3.4), and a case study (Section 3.5).

### 2.1 miRNA-gene and gene-gene interactions

We used the most comprehensive MGI dataset available in the literature. This dataset comprises 251,888,701 putative MGIs derived from miRNA target prediction algorithms and filtered using Argonaute (AGO) CLIP-seq data to retain experimentally supported miRNA–gene interactions (34). Specifically, 36 AGO-CLIP-seq datasets with 434,701 AGO-merged peaks from starBase v2.0 (35) were downloaded as the evidence of physical interactions between miRNAs and their target genes. To identify MGIs in 3’ untranslated regions (UTRs), MGIs downloaded from TargetScan v7.0 (36, 37) were intersected with AGO-merged peaks. For the identification of MGIs in coding sequence and 5’ UTRs, miRNA target sites predicted by RNA22 v2 (38) were first intersected with AGO-merged peaks, followed by validation using RNAhybrid v2.1.2 (39). Only the MGIs that were predicted as significant by both RNA22 and RNAhybrid and that overlapped with AGO sites were considered valid. The miRNA and gene names were unified using miRBase (40) and HGNC gene symbols v115 (updated 09/2025) (41). This resulted in a global MGI list containing 306,150 MGIs, 2,562 miRNAs, and 5,135 protein-coding genes.

We extracted 282,504 GGIs from Omnipath (42–44) and used them to reconstruct networks for pathways that have no predefined GGIs, such as cancer-specific and cancer hallmark pathways (Section 2.2). Only experimentally validated interactions for *Homo Sapiens* were retained, and undirected interactions were treated as bidirectional. For Reactome pathways, their corresponding networks were reconstructed using the database’s predefined GGIs (45). As a result, we obtained 464,595 GGIs involving 13,640 genes.

### 2.2 Network construction for biological pathways

We used three pathway datasets, including Reactome (45) and cancer hallmark pathways and cancer-specific pathways that were curated by us (Table S1). The public database Reactome contains 2,766 pathways (v94, updated 09/2025) comprising 11,426 unique genes. Of these, 3,122 are targeted by at least one miRNA in the global MGI list. We kept 2,394 pathways containing at least one MGI for our analysis. Cancer hallmarks are the tumors’ functional capabilities that are indispensable during malignant tumor development (46). We created cancer hallmark gene sets in our previous study (47), in which relevant gene ontology terms were curated into 10 cancer hallmarks comprising 6,637 unique genes, of which 1,734 are identified as miRNA targets in the global MGI list. We created cancer-specific pathways by searching for relevant keywords in MSigDB (v2025.1) (20) (Table S2). A total of 576 pathways were identified for 16 cancer types. Of those, 570 contained at least one MGI and were used as true positive pathways for benchmarking (Table S1). The Reactome and cancer-specific pathways contain gene sets ranging in size from 1 to 2,611 and 5 to 1,662, respectively. Cancer hallmark pathways have larger gene set sizes ranging from 538 to 3538. (Figure S2). After obtaining pathway gene sets, we constructed gene regulatory networks for pathways. We first mapped global MGIs onto pathway genes. We then incorporated GGIs, which are included when both genes appear in the pathway. Ultimately, we obtained pathway networks in which miRNAs and genes were the vertices and MGIs and GGIs were the edges.

### 2.3 Gene and miRNA expression data processing and analysis

Gene and miRNA expression profiles from both tumor and normal tissue samples were downloaded from The Cancer Genome Atlas (TCGA), with additional gene expression profiles for normal tissues retrieved from the Genotype-Tissue Expression (GTEx) project (48). We obtained read-count data for gene and miRNA expression profiles across 26 cancer types. The former comprises 8,768 tumor samples and 9,969 normal samples, while the latter comprises 8,630 tumor samples and 623 normal samples (Table S3). A total of 9,031 samples with paired miRNA and gene expression profiles (8,453 tumor samples and 578 normal samples) were used for downstream pairwise correlation analysis. To ensure sufficient statistical power, we excluded cancer types with fewer than three tumors or normal samples, or fewer than ten samples in total with paired miRNA and gene data. Consequently, we obtained data for 17 cancer types for further analysis.

We used DESeq2 in R to perform differential expression analysis for genes and miRNAs, respectively (49). We performed the analysis using genes and miRNAs with read counts greater than 10 in at least five samples. Differentially expressed genes (DEGs) and differentially expressed miRNAs (DEmiRs) were defined as those with Benjamini-Hochberg (BH) adjusted p-values ≤ 0.05 (50) and |log2 fold-change| ≥ 1 (tumor vs. normal samples). We performed Spearman correlation analysis using samples with both miRNA and gene expression profiles. Based on the assumption that effective miRNA downregulates the expression of target genes, MGIs exhibiting negative correlation coefficients and BH adjusted p-values ≤ 0.05 were recognized as effective interactions.

### 2.4 Node-based miRNA enrichment analysis algorithms

#### 2.4.1 Gene-centric methods

For node-based miRNA enrichment approaches, gene-centric methods identify a set of genes of interest by using the global MGIs list that links a list of miRNAs (e.g., DEmiRs) to their target genes. These genes are subsequently used as input data for enrichment analysis. In line with this, we implemented two gene-centric methods, including target gene over-representation analysis (TG-ORA) and target gene scoring-based analysis (TG-Score).

TG-ORA evaluates whether genes of interest are over-represented in a pathway using one-sided Fisher’s exact test. Under the null hypothesis that there is no significantly greater overlap between the interested genes and pathway genes than expected by chance, the p-value of enrichment is calculated using Equation (1).

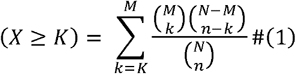

*n* represents the number of interested genes (e.g., target genes of DEmiRs). *K* represents the number of genes that overlap between the pathway genes and the interested gene list. *M* represents the number of pathway genes. *N* represents the total number of background genes (e.g., all genes in the pathways).

TG-Score performs enrichment analysis on a ranked list of genes and tests whether pathway genes are non-randomly concentrated toward the top or bottom of the ranked list. Specifically, by walking along the ranked gene list, the running-sum (RS) statistic increases when a gene hits in the pathway and decreases by a fixed value otherwise. The degree to which the RS increases is determined by the relative position of the gene *j* in the ranking list. A pathway’s RS is calculated using Equation (2).

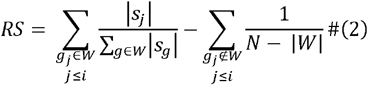

*s*_*j*_, *W*, |*W*|, *N* represent the ranking score of *j*_th_ gene, the pathway gene set, the pathway’s size indicated by the number of its involving genes, and the total number of genes in the ranked list, respectively. *j* ≤ *i* indicates the position of gene *j* up to position *i* in the ranking list, with *j* at the top or bottom of the list having bigger increase in RS than those in the middle of the list. The enrichment score (ES) is defined as the maximum deviation of the RS curve from zero. Genes are ranked by statistics that are genes’ fold-change divided by the corresponding standard error. A ES’s p-value is assessed through permutation test by randomly shuffling gene labels in the ranked list and recalculating ES values (51).

#### 2.4.2 MiRNA-centric methods

For miRNA-centric enrichment approaches, miRNAs are used to replace protein-coding genes in the analysis. Specifically, the pathway gene sets are converted into pathway-specific miRNA sets using the global MGI list. Like TG-ORA, miRNA over-representation analysis (MiR-ORA) is performed using miRNAs of interest (e.g., DEmiRs) as the input data (Equation 1). Like TG-Score, for miRNA scoring-based method (MiR-Score), miRNAs are ranked according to their log2 fold-change divided by the standard error and the enrichment scores are computed (Equation 2). Permutation tests are used to assess whether pathway-associated miRNAs are non-randomly enriched at the extremes of the ranked list.

### 2.5 Edge-based miRNA enrichment analysis algorithms

Conventional node-based methods treat genes and miRNAs as independent entities, thereby failing to capture the regulatory interdependencies encoded in MGIs and introducing bias into the pathway-level annotation of miRNA function. To address this limitation, we developed five edge-based enrichment methods that incorporate MGI data and network topology to analyze miRNA function. Firstly, we computed MGI scores to quantify the regulatory strengths of MGIs. MGIs that exhibit significantly negative correlations between DEmiRs and DEGs are defined as differentially expressed MGIs (DEMGIs) based on the following criteria.

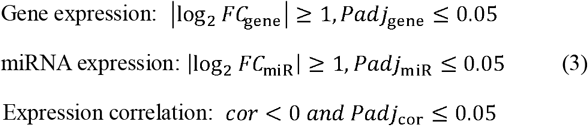

*cor* and *Padj*_*cor*_ represent the Spearman correlation coefficients and BH adjusted p-values that are computed using the data from tumor samples. log_2_*FC* and *Padj* represent log2 fold-changes and BH adjusted p-values that are obtained from differential expression analysis for miRNAs and genes. For each MGI, its relative expression change ratio and MGI score is defined in the following.

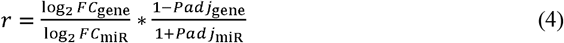

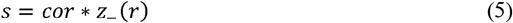

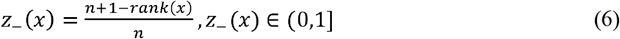

The ratio *r* captures the relative magnitude of gene expression change with respect to miRNA expression change, weighted by the statistical significance of differential expression analysis. Assuming that miRNAs repress gene expression (5, 6), a smaller negative value of *r* reflects a stronger repressive effect of a miRNA on its target gene. The MGI score *s* measures the strength of regulation considering expression correlation (*cor*) and relative expression change ratio (*r*). To mitigate the influence of extreme values and ensure comparable scaling of the two factors, we use the reverse rank-based function *z*_*-*_ to transform the ratio *r* (Equation 6). In *z*_*-*_, *n* represents the total number of *r*. Theoretically, the score *s* lies within the interval [-1, 1], where smaller values indicate stronger miRNA-mediated downregulation of genes and greater regulatory strength (Figure S3).

#### 2.5.1 Edge-ORA: MGI over-representation method

Inspired by TG-ORA, we developed edge-based over-representation method (Edge-ORA). Specifically, we used Equation (1) to test whether MGIs of interest are significantly over-represented in pathways. Pathways’ DEMGIs are the model’s input data, and the background list includes all MGIs identified in the pathways.

#### 2.5.2 Edge-Score: MGI scoring-based method

Inspired by TG-Score, we develop Edge-Score based using a ranked MGI list. Using Equations (4-6), all MGIs are ranked according to their MGI scores in descending order, where the highest positive score is first and the lowest negative score is last. Like Equation (2), the RS of a pathway is calculated as follows.

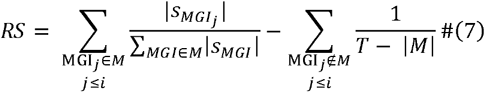

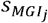 represents the *j*_th_ MGI score, *M* is the pathway’s MGI set, |*M*| is the MGI size in a pathway, and *T* is the total number of MGIs in the ranked list. A pathway’s ES is defined as the computed RS curve’s maximum deviation from zero from the negative side. We considered negative ESs to be biologically significant since MGI with negative scores contain effective miRNAs for downregulating gene expression. We applied permutation test to obtain the ESs’ p-values (51). Notably, because the more negative MGI scores indicate stronger gene regulation by miRNAs, only pathways with negative-sided significance are considered enriched.

#### 2.5.3 Edge-2Ddist: MGI two-dimensional scoring method

Because the expression correlation and expression change ratio of miRNAs and their target genes can indicate the regulatory strength of MGI (Equation 4), we developed the Edge-2Ddist method. This method uses the distribution of the two variables on a two-dimensional (2D) coordinate for enrichment analysis. Similar 2D distribution analysis have also been proposed by others (52, 53). Specifically, MGIs are mapped into a 2D coordinate space, where the x-axis represents the negatively normalized Spearman correlation coefficient 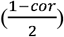 and the y-axis represents the normalized expression change ratio (*z (r)*; Equation 6). Both x and y values are scaled to the range of [0, 1], where higher values along each axis indicate stronger gene regulation by miRNAs. MGIs located along the diagnosis line up to the upper-right corner of the coordinate space corresponds to stronger gene regulation by miRNAs. This is characterized by smaller negative correlation and higher relative expression change ratio. The score (*S*_*i*_) for pathway *i* is calculated as the sum of the distances of its MGIs to the null point (Equation 8), where the *cor*_*ij*_ and *r*_*ij*_ are the corresponding values of MGI *j* in pathway *i*, and *n*_*i*_ is the total number of MGIs in pathway *i*.

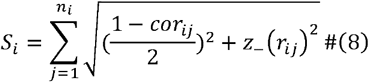

To evaluate a pathway’s statistical significance, we performed permutation tests by randomly selecting the same number of MGIs from the global MGI list to form random pathways and calculating their pathway scores. We repeated the process 1,000 times. Since MGIs’ distances to the null point are positive values and larger scores reflect stronger miRNA regulatory strength, we assessed pathway significance using a one-sided test in the positive direction. To handle the extremely small, estimated p-values obtained using the permutation test, we used the z-score method to normalize all pathway scores (*S*) using random pathway scores (*S*_*r*_) (Equation 9).

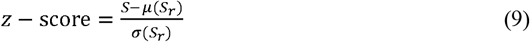

*μ*(*S*_*r*_) and *σ*(*S*_*r*_) correspond to the average and the standard deviation of the 1,000 random pathways’ scores that follows a Gaussian distribution *N* (0,1). The positive-sided p-value is approximated via Taylor expansion of the distribution integral, where larger z-scores correspond to stronger aggregate gene regulation by miRNAs within a pathway.

#### 2.5.4 Edge-Topology: MGI pathway network method

Inspired by a previous study (21), we developed the Edge-Topology method by incorporating the topology of pathway networks into MGI-based enrichment analysis. For each pathway, we used its containing MGIs and GGIs to construct a directed network. For pathway *i*, its score *S*_*i*_ is defined using MGIs’ scores (Equation 5) and edge betweennesses in the pathway:

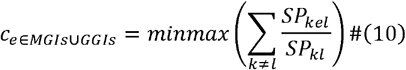

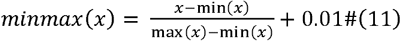

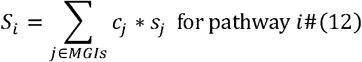

Here, *c*_e_ denotes the min-max normalized edge betweenness centrality of the edge *e*, defined as the sum over all node pairs (*k, l*) of the fraction of shortest paths between *k* and *l* that pass through edge *e*. Specifically, *SP*_*kl*_ represents the total number of shortest paths between vertices *k* and *l* while *SP*_*kel*_ represents the number of those shortest paths that pass through edge *e*. The raw edge betweenness values are normalized using the min-max transformation in Equation (11). To prevent MGI information loss arising from edges with a betweenness centrality of zero, we added a constant of 0.01 to all edge values prior to min-max normalization (Figure S4). Consequently, for each MGI, its corresponding centrality score *c*_*j*_ is a normalized edge betweenness value in the pathway network, and its MGI score *s*_*j*_ is calculated as defined in Equation (5). The pathway score *S*_*i*_ therefore presents a weighted sum of MGI scores, where weights reflect the topological importance of MGIs within the pathway. We assessed statistical significance of each pathway score using a permutation test (n = 1,000). In each permutation, target genes within the pathway network were randomly reassigned to the pathway miRNAs while preserving the node degrees of both miRNAs and genes, with GGIs held unchanged. MGI and centrality scores for the permuted interactions were computed using Equations (4–6) and (10–11), respectively, and the corresponding pathway score was derived using Equation (12). Since MGIs with smaller negative scores and higher centrality scores are more important, pathway significance was evaluated using z-scores as defined in Equation (9), with p-values estimated from the negative-sided tail of the permutation distribution.

#### 2.5.5 Edge-Network: MGI global network method

Inspired by NetPEA (22), we developed Edge-Network that uses random walk with restart (RWR) on a gene regulatory network to identify enriched pathways by diffusing regulatory signals from MGIs of interest (e.g., DEMGIs). For each pathway source (Reactome, cancer-specific, and hallmark pathways), we constructed a directed network using the corresponding MGIs and GGIs (Table S3). Within each network, DEMGIs are designated as seed MGIs, assigned initial edge weights from which flux propagation originates. We then applied the RWR algorithm to propagate seed MGI fluxes through the network according to the rule defined in Equation (13).

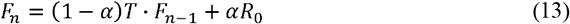

The restart probability *α* determines the percentage of edge fluxes (*F*) retained to itself at integration *n*. We set *α* = 0.5, such that at each iteration, 50% of the flux in an upstream edge evenly propagates to its downstream edges that share a common node with it. The restart vector *R*_*0*_ is the initial edge weights. Seed MGIs’ edge weights are set to 1 and the other edges are set to 0. The transition matrix T governs the diffusion of edge flux through the network. Specifically, flux propagates from an upstream edge to its downstream edges via shared nodes, defined as the target node of the upstream edge and the source node of the downstream edge. For GGIs having no downstream edges, self-loops were added to retain their flux and prevent the total edge flux from converging to zero over successive iterations. For isolated seed MGIs that are not connected to any other edges, flux diffusion follows the same rule defined in Equation (13), resulting in a gradual decrease in flux over iterations. The algorithm terminates when either the sum of flux changes across all edges falls below a predefined tolerance of 10□, or the number of iterations reaches the maximum threshold of 100. Afterwards, the enrichment score (*S*_*i*_) for pathway *i* is the sum of all MGIs’ and GGIs’ final edge fluxes (Equation 14).

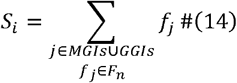

The statistical significance of each pathway score is assessed via permutation testing to determine whether the observed pathway accumulates more edge flux than expected by chance. During permutation, the network structure is preserved while the restart vector *R*_*0*_ is randomly resampled. Specifically, isolated seed MGIs are assigned an edge flux of 1 and all GGIs an edge flux of 0. An equal number of connected seed MGIs are randomly reassigned among all connected MGIs with their edge fluxes set to 1, while the remaining MGIs are assigned an edge flux of 0. Permuted pathway scores *S*_*r*_ are computed by repeating the (RWR) algorithm across 1,000 permutations. Since edge fluxes are non-negative and higher values reflect stronger aggregate effects of the seed MGIs, pathway significance is evaluated using z-scores and one-sided p-value estimation in the positive direction, as defined in Equation (9).

For all nine methods in miREA, we applied multiple testing corrections to control false discovery rates (54). We considered pathways with BH adjusted p-values ≤ 0.05 significantly enriched.

### 2.6 Methods’ ability to identify cancer-associated miRNAs and genes

To test whether our methods can identify miRNAs and genes associated with cancers, we performed analyses of 17 cancer types for which there are sufficient samples that meet the minimum requirements for statistical testing (Table S3). Specifically, we compared the overlap of the important genes and miRNAs identified by four methods with the curated cancer-associated genes and miRNAs from three databases, including 139 miRNAs from Cancer miRNA Census (CMC) (55), 1,200 genes from oncoKB (v24, Nov 2025) (56, 57), and 586 tier-1 cancer genes from COSMIC (v103, 18 Nov 2025) (58) (Table S4). TG-Score and Edge-Score identify important genes or MGIs appearing in the ranked list up to the point at which the RS statistic reaches its maximum deviation from zero. For Edge-ORA, important MGIs are defined as DEMGIs occurring in statistically enriched pathways (BH-adjusted p-value ≤ 0.05). Edge-Network expands the DEMGI list by incorporating MGIs and GGIs with non-zero edge flux following RWR simulation. From these method-specific interaction sets, we derived lists of important miRNAs and genes were subsequently derived. For TG-Score, important miRNAs were extracted from the global MGI list based on the identified important genes. For all edge-based methods, important miRNAs and genes were extracted directly from the identified important MGIs. Notably, the lists generated by Edge-Network are considerably larger, as the method operates on a network integrating all molecular interactions within a given pathway resource. The overlap between these lists and known cancer-associated miRNAs and genes from the three reference databases was then quantified using the geometric mean of two proportions: the fraction of identified important miRNAs or genes that are annotated as cancer-related (*p*_1_), and the fraction of known cancer-related miRNAs or genes that are recovered as important ones (*p*_*2*_). A higher geometric mean indicates greater consistency with existing biomedical knowledge, and therefore better biological interpretability.

### 2.7 Strategy and parallel computing for efficient computation

Given the integration of multi-omics data and large-scale networks, the edge-based methods are more computationally demanding than conventional node-based approaches. We implemented several optimization strategies to improve computational efficiency. For Edge-2Ddist, pathway-level parallelization was applied to compute pathway scores and assess statistical significance simultaneously. For Edge-Topology, pathways were distributed across CPU cores with approximately balanced total pathway sizes per core, and an early-stopping strategy was employed to terminate permutation testing for non-significant pathways. Specifically, an initial permutation of 100 iterations was performed; pathways for which more than 50 permuted scores exceeded the observed score were excluded. The remaining pathways underwent an intermediate round of 500 permutations, with the same exclusion criterion applied. Only pathways with strong evidence of enrichment proceeded to the full permutation test (n = 1,000). This approach substantially reduces computational overhead while preserving statistical power. For Edge-Network, parallel computing was applied to all pathways during the permutation test. Each iteration included restart vector resampling, RWR, and pathway score computation. These steps were executed independently across CPU cores.

### 2.8 Reproducibility

We developed miREA using R v4.4.2. We run all computations on the Finnish CSC-IT Center for Science computer cluster. The GitHub repository contains documentation describing all the packages, data, and codes used to reproduce the results (https://github.com/laixn/miREA).

## 3 RESULTS

### 3.1 miREA contains five edge-based methods for miRNA-oriented enrichment analysis

We develop five miRNA-oriented enrichment analysis methods that use MGI edges and network topology to analyze miRNA’s functions at the pathway level (Figure 1 and Table S5). Edge-ORA uses DEMGIs (i.e., MGIs with differentially expressed miRNAs and target genes that exhibit significantly negative correlations, as defined in Equation (3) as input data. Then, it performs hypergeometric tests to determine if the proportion of DEMGIs in a pathway is higher than expected by chance. We further develop Edge-Network by combining DEMGIs and a pathway network which contains MGIs and GGIs across all pathways of interest. We set DEMGIs as seed MGIs in the network and run RWR algorithm to simulate the diffusion of edge fluxes over the whole network. The simulations indicate how the DEMGIs could affect their downstream edges, and test whether a pathway’s edges accumulate more diffused fluxes compared to random selection of seed MGIs. Furthermore, we develop three methods, namely Edge-Score, Edge-2Ddist, and Edge-Topology, by utilizing MGIs’ scores (Equations 4-6). The scores contain two terms: correlation coefficients between miRNAs and their target genes, and the expression ratio between the expression fold-change of miRNAs and their target genes. The former indicates how the expressions of both molecules are correlated, and the latter indicates the extent to which changes in the expression of miRNAs can affect the expression of their target genes. As miRNAs should downregulate gene expression except in a few cases (5, 6), we expect a stronger MGI with smaller negative correlation coefficient and larger ratio. Inspired by previous studies (52, 53), we develop Edge-2Ddist that analyze MGIs distributed on a 2D coordinate, for the normalized expression correlation and the expression ratio. We compute a score for each pathway by summing the distances of its containing MGIs to the null point in the coordinate system and use permutation test to determine whether the pathway’s MGIs are statistically different from random MGIs in the 2D coordinate system. We further transform MGIs’ correlation and ratio values scores into one score to develop Edge-Score that resembles the idea of the most widely used tool gene set enrichment analysis (20). The method ranks MGIs and computes enrichment scores for pathways followed by permutation tests for getting p-values based on their scores. Finally, we incorporate pathway networks and MGI scores to develop Edge-Topology. This method maps MGI scores onto network edges and combines edge betweenness coefficients with the score to identify enriched pathways. We compute the results’ significance using permutation test.

**Figure 1.**
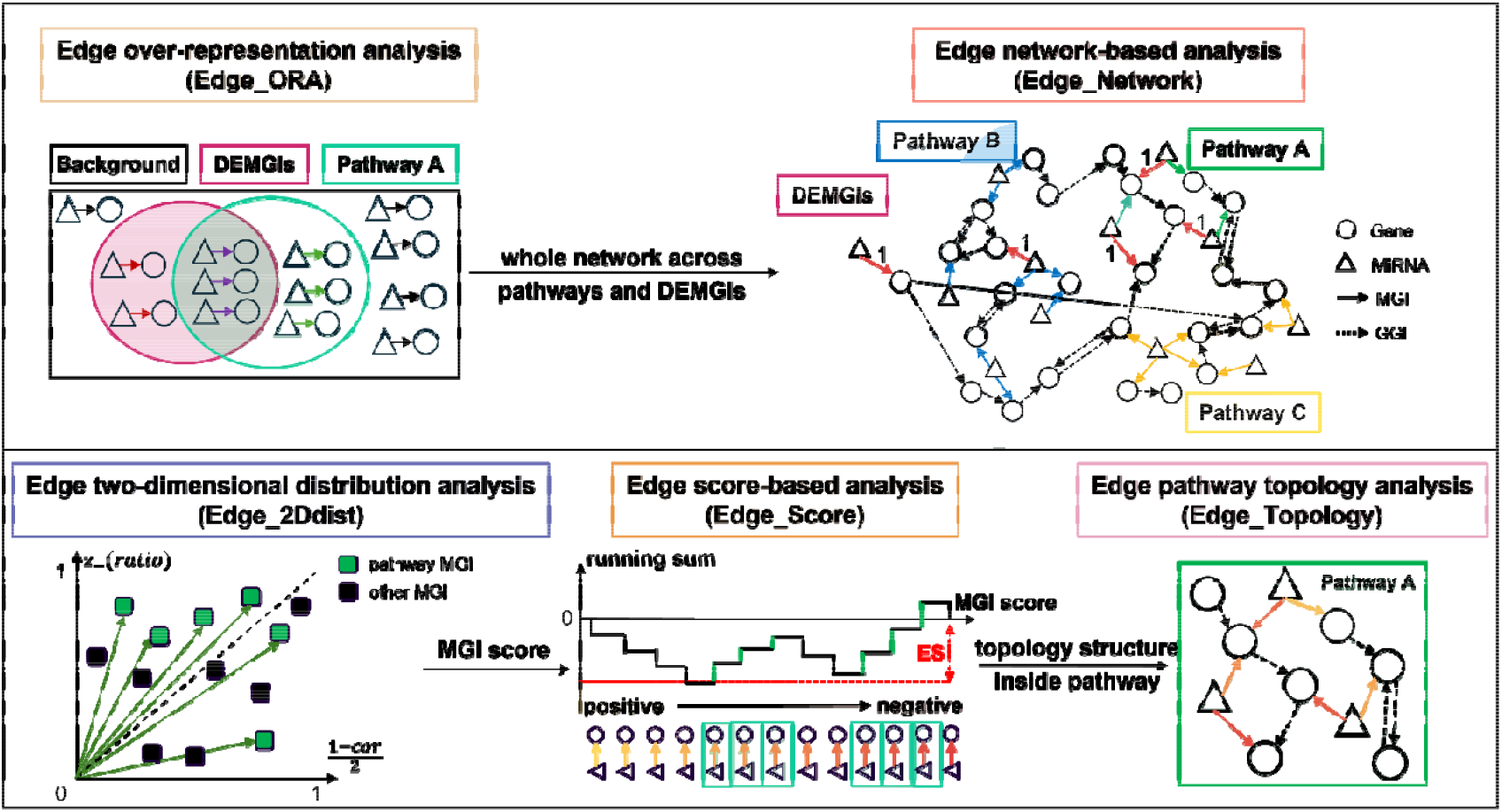
Overview of the five edge-based enrichment analysis methods. Edge-ORA applies hypergeometric tests on a list of MGIs of interest to identify enriched gene sets from public databases or curated pathways. Edge-Network extends the framework by considering pathways’ network topology using the RWR algorithm and permutation test. Furthermore, we consider quantitative information in MGI and develop three methods. Edge-2Ddist consider two information in MGIs including correlation and relative expression change ratio between miRNAs and their target genes. Edge-Score transforms the two pieces of information into an MGI score. Based on the MGI score, Edge-Topology further incorporates the pathway network structure. Each part illustrates an edge-based method, while the arrows and text in between indicate the progressive information introduced among different methods. Triangles and circles represent miRNAs and genes, respectively. Arrows indicate the direction of regulation, with solid lines denoting MGIs and dashed lines denoting GGIs. In the Edge-2Ddist, squares represent MGIs. For example, pathway A contains seven MGIs, two of which are DEMGIs, and six GGIs. The MGIs involved in pathway A are highlighted with green shading or arrows. Similarly, pathway B and pathway C in the Edge-Network are depicted in blue and yellow shading. DEMGIs are indicated by pink shading and pink arrows. We visualize MGI scores via arrows with a yellow-to-red gradient, representing the transition from positive to negative scores. The enrichment score (ES) is the curve’s maximum negative deviation from zero.

### 3.2 Evaluation of the miREA methods’ performance

#### 3.2.1 The miREA methods achieve improved true positive rates with controlled false positive rates

Using cancer-specific pathways that are curated from MSigDB and regarded as true positives (TPs), we benchmark four node-based and five edge-based methods in 16 cancer types (Methods Section 2.2). For each cancer type, we compute true positive rate (TPR) that is the percentage of enriched TP pathways (adjusted p-values < 0.05) for the positive benchmark. Notably, the cancer-specific TPRs are influenced by the number of cancer-specific pathways, we further calculate the overall TPR as the total number of enriched TP pathways divided by the total number of TP pathways across all cancer types. To create negative pathways, we randomly shuffle the MGI’s data used in the cancer-specific TP pathways while preserving their topologies. Consequently, the resulting expression profiles and edge profiles lack biological meaning in the negative pathways. For each cancer type, we shuffle their differential expression analysis results for miRNA and their target genes. Specifically, for TG-ORA, TG-Score, MiR-ORA, MiR-Score, and Edge-Topology, we randomly assign entries to miRNAs and genes. For Edge-ORA, Edge-Score, Edge-2Ddist, and Edge-Network, we shuffle MGI scores by randomly assigning the entries to MGIs. To ensure sufficient statistical power and obtain an enlarged number of negative pathways, we repeat the randomization process 30 times using the 576 TP pathways, resulting in 17,100 pathways (570×30 times) in total (Table S2) (59). Finally, we perform enrichment analysis on the negative pathways and compute the false positive rate (FPR) which is the percentage of pathways to have p-value < 0.05.

Among the node-based methods, TG-Score is the only method that can identify cancer-specific pathways with the overall TPR of 0.142, while the other three fail to detect biologically meaningful pathways (Figure 2A). This is likely due to the dilution of direct biological associations during the many-to-many mapping process between miRNAs and their target genes. In contrast, edge-based methods have better performance in identifying cancer-specific pathways, indicating the methods’ greater sensitivity in identifying biologically relevant pathways. Specifically, Edge-Score achieves the highest TPRs in 9 out of 16 cancer types and has the best overall TPR (0.267). However, Edge-Topology’s performance is poor, which may be due to the method’s conservative assumption. Compared to Edge-Score, which requires pathway MGIs to exhibit strong regulatory strengths for enrichment, Edge-Topology is much more conservative. Edge-Topology requires that MGIs within the pathway exhibit both strong regulatory strengths and high topological importance, as indicated by edge betweenness centrality. Using this method, Reactome and cancer-specific pathways with smaller pathway sizes get larger p-values than cancer hallmark pathways (Figure S5), which is consistent with studies that topology-aware methods are more sensitive to the selection of pathway databases (60, 61). This reflects the limited network connectivity of smaller pathways, which reduces the degree of randomness during permutation testing. Conversely, larger pathways with denser interaction networks provide a bigger permutation space, yielding more stable null distributions and, consequently, more statistically robust enrichment results. Furthermore, Edge-Network integrates all pathway sets into one network, increasing the robustness of enrichment analysis by leveraging global network topology. However, propagation across a large network may dilute the contribution of individual MGIs. This renders the method less sensitive to pathway-specific signals, which could compromise its ability to detect enrichment in smaller or less densely connected pathways.

**Figure 2.**
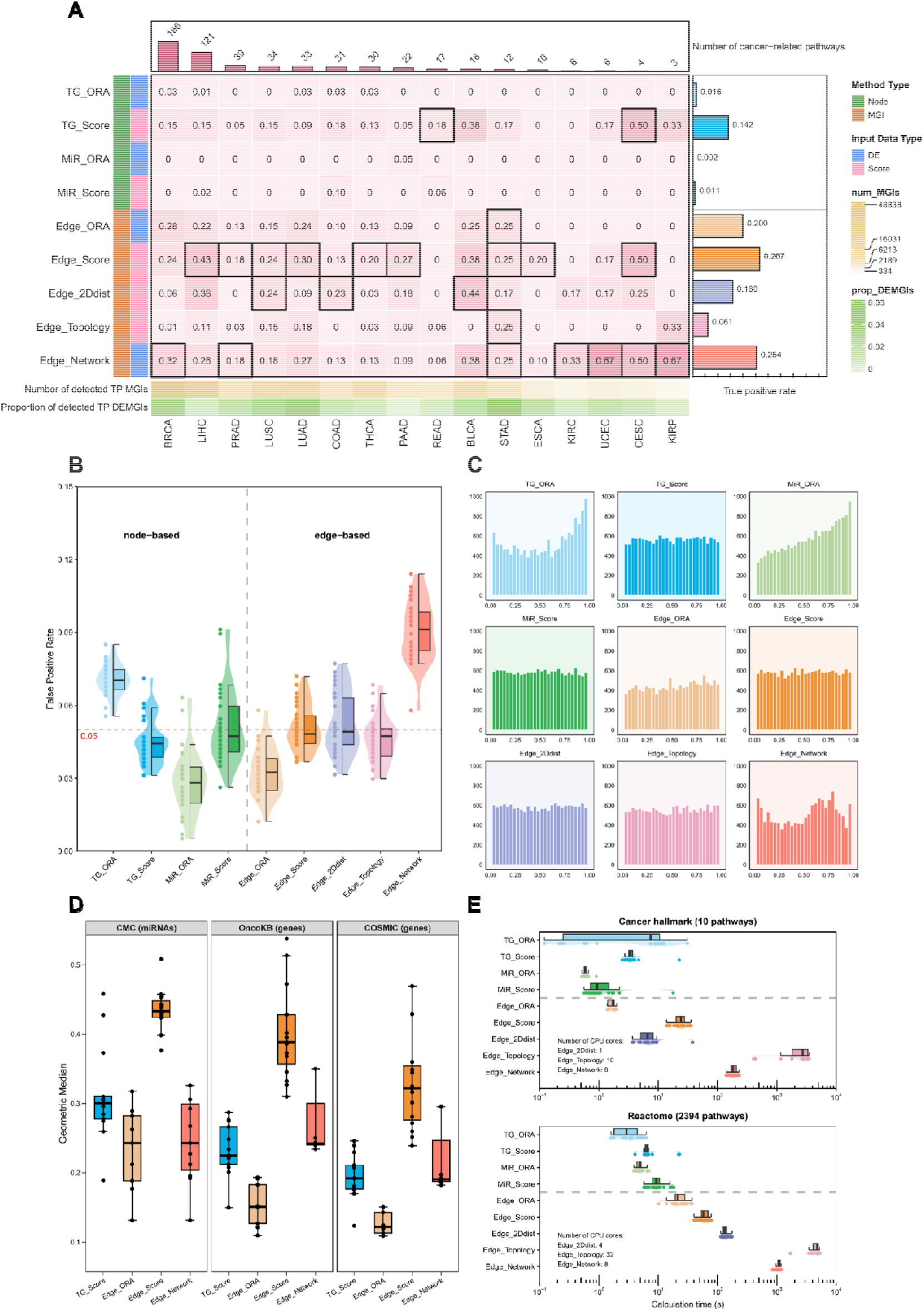
Benchmark performance for nine enrichment analysis methods. **(A)** The heatmap shows the nine methods’ (per row) TPR for positive pathways in 16 cancer types (per columns). The grids in black frames highlight the best-performing method with the highest TPR for a cancer type. The top bar shows the total number of identified cancer-related pathways for cancers. The right bar plot shows the methods’ overall TPRs, which is the ratio of the total number of enriched pathways across all cancer types to the total number of TP pathways. The bottom annotation shows the number of MGIs and the proportion of DEMGIs in in all curated pathways of a cancer type. **(B)** The boxplot summarizes the distribution of FPRs for each method, where the bottom and the top of the box indicate the 25% and 75% quantile, and the black line in the middle indicates the median FPR across 30 randomizations. Each dot corresponds to an overall FPR for one randomization. The red dashed line indicates the predefined nominal significance level of 0.05 **(C)** The histogram illustrates the distribution of negative pathways’ p-values for each method. **(D)** Overlap between important miRNAs or genes and annotated cancer-related miRNAs and genes in cancer hallmark pathways. Each dot represents one cancer type. The number of dots may be smaller than the number of cancer types because no enriched pathways are identified in some of them. **(E)** Summary of calculation times for different methods, including enrichment for cancer hallmark (upper) and Reactome (lower) pathways. Each dot represents one cancer type. The methods that use multiple CPU cores for computing are highlighted otherwise the computation is done using a single core.

For the negative pathways, a good method should have no preference in p-value distribution. Thus, we expect that unbiased methods should have an approximately uniform p-value distribution for all negative pathways, resulting in a proportion of pathways with p-values below a predefined nominal significance level (*α* = 0.05). Our edge-based methods show good control of FPR (Figure 2B), with Edge-Score, Edge-2Ddist, and Edge-Topology showing median FPRs close to 0.05. Edge-ORA maintains stricter with a median FPR of approximately 0.03, while Edge-Network exhibits less stringent control with a median FPR of around 0.09.

We further investigate the p-value distributions generated in the true negative data set (Figure 2C). Most methods remain neutral and generate approximately uniform p-value distribution. TG-ORA and MiR-ORA produce p-values that are skewed toward 1, and Edge-Network exhibits mild irregular deviations from uniformity. In addition, we compare p-value distributions between true positive and true negative pathways (Figure S6). All methods except TG-ORA and MiR-Score exhibit significantly different distributions (Fisher-Pitman p-value smaller than 0.05) between two pathway data, suggesting that edge-based methods’ discriminative ability in distinguishing positive pathways from negative ones. In addition, because negative-sided MGI scores are considered to have strong regulatory strengths, we determine enrichment based on one-sided p-value for edge-based methods, which exhibit bimodal p-value distribution for TP pathways.

#### 3.2.2 The miREA methods can distinguish pathways for specific cancer types

To evaluate the ability of our methods to correctly identify pathways specific to a cancer type among all 570 TP pathways, we conduct enrichment analysis on 16 cancer types (Figure S7). For each cancer type, we consider its corresponding TP pathways as true pathways, and other TP pathways that belong to other cancer types as false pathways (Methods Section 2.2). In each cancer, we rank pathways based on their enrichment p-values, resulting in a rank score from 0 to 1. Pathways with smaller p-values have smaller rank scores. Then, we use the median rank of each cancer type’s true pathways to compare methods’ performances. The expectation is that a better method will have a lower median rank score for true pathways. Except Edge-Topology, our methods are among the top-performing ones to distinguish cancer-specific true pathways from the false ones (Figure S7, left). Additionally, we aggregate all TP pathways’ ranks from all cancer types and compare the rank of the true and false pathway groups for each method. We expect that a good method should show lower scores for the pathways in the true group than in the false group. As with the median rank comparison, methods with better performance show clear differences between the true and false groups. However, TG-ORA, Edge-2Ddist, and Edge-Topology show no significant differences for both groups (Figure S7, right). MiR-ORA and MiR-Score show even higher scores for pathways in the true group. Taken together, TG-Score, Edge-ORA, Edge-Score, and Edge-Network show comparable ability to identify specific cancer-related pathways than other methods.

#### 3.2.3 The miREA methods can identify miRNAs and genes associated with cancers

We further compare methods’ ability to identify miRNAs and genes associated with specific cancers, and methods with better performance can provide more accurate identification of cancer-specific regulatory mechanisms. We focus on four methods (i.e. TG-Score, Edge-ORA, Edge-Score, and Edge-Network) that have the best performance at the pathway-level and use their identified important genes or MGIs in 10 cancer hallmark pathways in 17 cancer types for the analysis (Methods Section 2.6). Compared to the best-performing node-based method TG-Score, Edge-Score is better at identifying important miRNAs and genes, Edge-Network performs similarly for genes but worse for miRNAs, and Edge-ORA performs worse for both miRNAs and genes (Figure 2D). This can be attributed to methodological differences. Edge-ORA relies solely on DEMGIs, Edge-Network uses both DEMGIs and network topology, and Edge-Score uses MGI scores to transform the information of DEMGIs into regulatory interaction strength. By incorporating more information, these methods achieve greater specificity in identifying experimentally validated cancer-associated miRNAs and genes.

### 3.3 Systematic analysis of the miREA methods

#### 3.3.1 The edge-based methods are robust

We further assess the robustness of the five edge-based methods. Among them, Edge-Score, Edge-2Ddist, and Edge-Topology use quantitative MGIs (i.e., MGI scores), while Edge-ORA and Edge-Network use qualitative MGIs (i.e., DEMGIs). For Edge-Score, Edge-2Ddist, and Edge-Topology, we evaluate whether the MGI scores’ variability can influence the methods’ FPR for 17,100 negative pathways. These pathways are biologically meaningless because they have the same topology as 570 TP pathways from 16 cancer types but contain randomly shuffled MGIs data that is repeated 30 times. Specifically, we perform Pearson correlation analyses and examine the influence of MGI scores’ standard deviation, value range, interquartile range, and median absolute deviation on the methods’ FPR. The insignificant correlations between the MGI scores’ profiles and the methods’ FPR indicate that the methods’ performance are robust to the variability of MGI scores (Figure S8).

For Edge-ORA and Edge-Network, we evaluate the effect of the number of the DEMGIs determined by log2 fold-change and adjusted p-value on the analysis results. We use the 10 cancer hallmark pathways and the 2,394 Reactome pathways instead of cancer-specific pathways. This allows for a fair comparison because the analysis uses the same pathways for each cancer type. The results show that relaxing both thresholds increases the number of DEMGI leads and results in higher overall positive rates for both methods (Figure S9). This suggests that a greater number of DEMGIs identified from data sets can provide more gene regulatory information, thereby increasing the methods’ sensitivity to identify relevant pathways.

#### 3.3.2 Integrating miRNA and gene expression data can improve the models’ performance

The methods in miREA can be categorized based on their required data sources for enrichment analysis. MiR-ORA, TG-ORA, and MiR-Score require miRNA expression data only, as their input data are derived from DEmiRs, DEmiRs’ target genes, or a ranked miRNA list based on expression levels, respectively. Edge-ORA and Edge-Network require DEMGIs, which can be derived from either miRNA expression alone or both miRNA and gene expression data. TG-Score, Edge-Score, Edge-2Ddist, and Edge-Topology require both miRNA and gene expression data. Here, we evaluate the performance of Edge-ORA and Edge-Network with and without gene expression profiles by performing enrichment analyses using either miRNA expression data alone or both miRNA and gene expression data. When using miRNA expression data alone, Edge-ORA and Edge-Network outperform other node-centric methods that require only miRNA data in identifying true positive pathways (Figure S10). Furthermore, incorporating DEMGIs derived from both miRNA and gene expression data further improves the performance of both methods. These results demonstrate that while edge-based methods are inherently miRNA-oriented, the availability of gene expression profiles can substantially enhance their performance.

#### 3.3.3 Parallel computation for improving the efficacy of miREA

We implement parallel computing and evaluate computational efficiency using different numbers of CPU cores on the bladder urothelial carcinoma (BLCA) dataset with two pathway sets. The first set comprises 10 cancer hallmark pathways with gene set sizes ranging from 538 to 3,538. The second comprised 2,394 Reactome pathways with gene set sizes ranging from 1 to 2,611, the majority of which are small gene sets (Figure S2). Although parallel computing provides limited benefits for Edge-2Ddist, it dramatically reduces the time required for Edge-Topology and Edge-Network, which require heavy computation of network topology (Figure S11). Based on the analysis, we select the suitable number of cores for specific methods and run the enrichment analysis across 17 cancer types using the cancer hallmark and Reactome pathways. The analysis with node-based methods can be accomplished on average in about 30 seconds (Figure 2E). Due to the use of multiple data sources and complex algorithms, analysis with edge-based methods requires on average 1 second to 1.5 hours. The parallel computing reduces their runtime to an acceptable level for computers with an 8-core CPU. Edge-Topology requires the longest computation time, reaching a maximum of ∼59 and ∼89 minutes for the cancer hallmark and Reactome pathways, respectively. The other methods complete the same analysis within ∼4 minutes and ∼20 minutes.

### 3.4 A summary of the miREA methods’ performance

We summarize the quantitative evaluation results across cancer types, as discussed in the previous two sections. These results include: (i) sensitivity, as indicated by TPR; (ii) specificity, as indicated by FPR; (iii) distinguishability, as indicated by the median rank for distinguishing cancer-specific pathways from other cancer-related pathways; (iv) biological interpretability of miRNAs and genes, as indicated by the geometric median of overlap between important miRNAs or genes and known cancer-related miRNAs and genes in cancer hallmark pathways; and (v) calculation efficiency, as indicated by the computational time for cancer hallmark and Reactome pathways. (Figure 3). Overall, compared to the node-based methods, the edge-based methods show better sensitivity and comparable specificity, distinguishability for pathways, and interpretability for miRNAs and genes. However, since some edge-based methods such as Edge-Topology and Edge-Network require the computation of network topology, their computation efficacy is quite lower than the node-based methods. Among the edge-based methods, Edge-ORA shows good sensitivity, specificity, and ability to distinguish pathways. However, it provides limited ability to interpret biological information for identifying consensus cancer genes and miRNAs. Edge-Score shows balanced sensitivity and specificity, as well as the best interpretability. Edge-2Ddist has compromised sensitivity, good control of false positives, and the worst distinguishability. Edge-Topology is the most conservative and time-consuming approach. It is characterized by the lowest sensitivity, the longest computation time, and distinguishability comparable to that of the other edge-based methods. Its poor performance in identifying pathways could be attributed to the long-tail distribution of the network’s edge betweenness, which results in a few MGIs dominating and most MGIs’ contributions being undifferentiable. (Figure S4). Edge-Network achieves the second-highest sensitivity, good distinguishability, and the second-best interpretability, but has compromised specificity.

**Figure 3.**
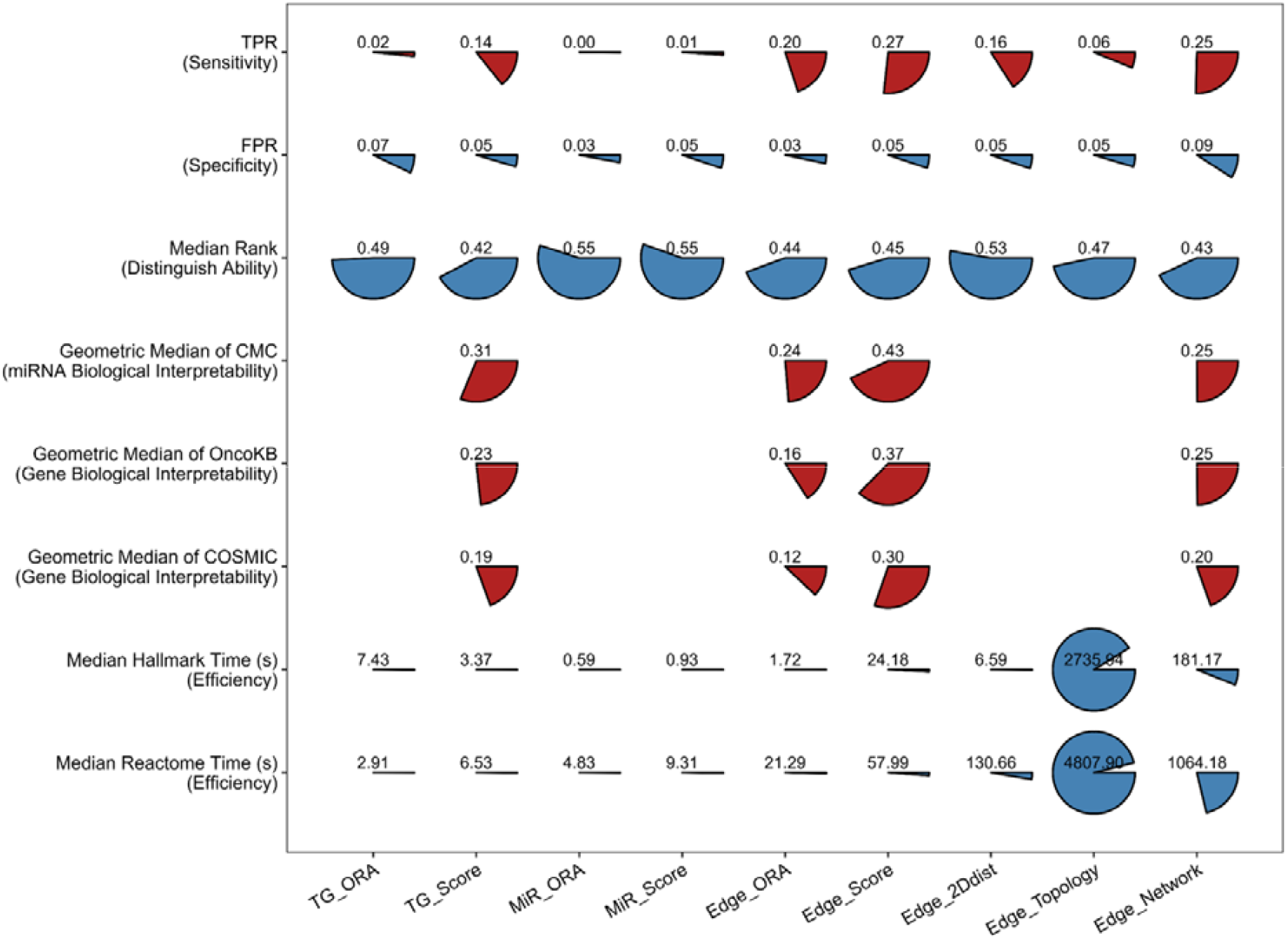
Performance metrics of methods. The pie charts show the values of different evaluation metrics for nine different methods Each row represents an evaluation metric, and each column represents a method. The metrics with higher values indicating better performances are filled in red. Conversely, those with lower values indicating better performances are filled in blue. Units corresponding to methods that do not generate current metric are left in blank. The TPR, FPR, geometric median of CMC, OncoKB, and COSMIC are overall values across all cancer types. The median rank, median hallmark time, and median Reactome time represent the median value of median relative ranks (Figure S7, left), calculation time of cancer hallmarks (Figure 2E, upper), calculation time of Reactome pathways (Figure 2E, lower) for different cancer types, respectively. The data used for the plot can be found in Table S6.

### 3.5 Characterizing miRNA-gene-pathway regulatory mechanisms in bladder cancer

To demonstrate the power of miREA in biomedical applications, we present a case study using the bladder urothelial carcinoma (BLCA) data from TCGA and GTEx (Table S7). Bladder cancer ranks as the ninth most common cancer worldwide, with an estimated 614,000 new cases and 220,000 deaths reported in 2022 (62), and shows the highest incidence rates in southern and northern Europe. Urothelial carcinoma is the most frequently observed subtype, accounting for approximately 90-95% of all bladder cancer cases (63).

We perform enrichment analysis using miREA’s nine methods across Reactome, cancer-specific, and cancer hallmark pathways (the full results can be found in the Supplementary Files 2-4). Here, we focus on analyzing and discussing the cancer hallmark pathways, as they represent consensus cancer’s capabilities that facilitate biological interpretation and method comparison. Six of the nine methods identify at least one significantly enriched pathway among the 10 tested (Figure 4A). MiR-ORA, MiR-Score, and Edge-Topology identify no enriched pathways, whereas Edge-Score and Edge-2Ddist identify the most. Pathways identified by methods within the same category (node-based or edge-based) show greater overlap than those identified across categories. Among the edge-based methods, Edge-Network demonstrates the highest concordance with other methods, as reflected by both pathway overlap analysis (Figure 4B) and principal component analysis (Figure 4C). Similar results are obtained using Reactome and cancer-specific pathways (Figure S12). Given its ability to capture consensus regulatory patterns across methods, we select Edge-Network to further characterize miRNA regulatory mechanisms in BLCA (Figure 4D). The top 10 miRNAs with the largest numbers of DEMGIs are enriched in five cancer hallmark pathways, namely sustained angiogenesis, evading growth suppressors, tissue invasion and metastasis, tumor-promoting inflammation, and evading immune destruction. Five of the top miRNAs are cancer-related as reported in the CMC (55), with the supporting evidence that some of their target genes are also cancer-related according to COSMIC (58) and OncoKB (56). The top miRNAs target 52 genes, 49 of which are upregulated and three of which are downregulated in the pathways. This results in 71 DEMGIs that contain 33 genes in enriched cancer hallmarks. We think that these MGIs are likely to be observed in experiments because the corresponding miRNAs and their target genes have differential expression changes and significantly negative correlations.

**Figure 4.**
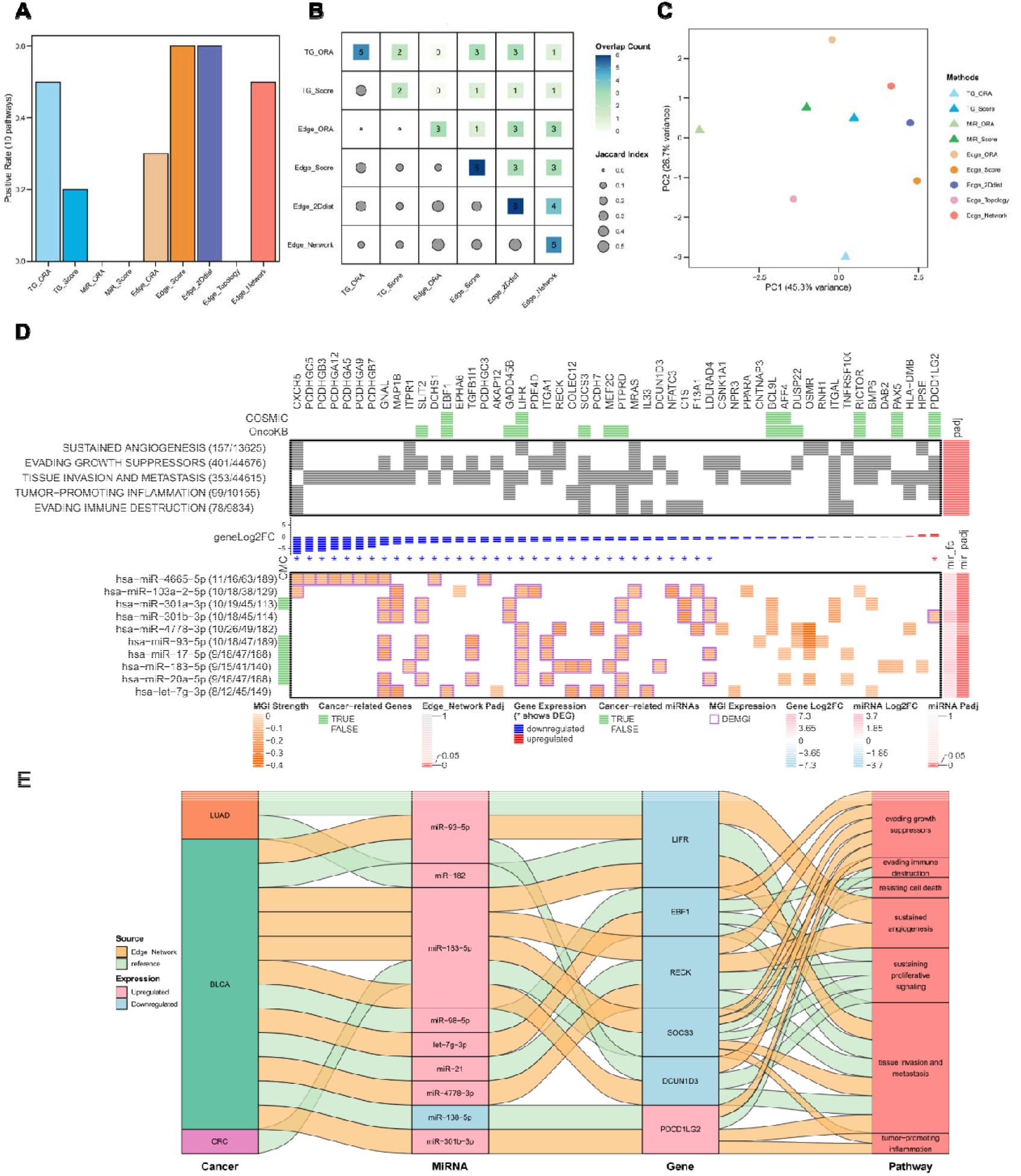
The BLCA case study. **(A)** The bar plot shows the identified enriched cancer hallmark pathways by nine different methods in miREA. (**B**) The plot shows the similarity of the identified pathways by different methods where the upper and lower triangles show the number and the Jaccard index of overlapped enriched pathways, respectively. Only methods with the identified enriched pathways are visualized. (**C**) The PCA analysis shows the clustering of methods based on the computed p-values for all 10 pathways for each method. The data used to create the figure panels A-C can be found in Supplementary Excel File 2. **(D)** The heat map shows the identified MGIs using Edge-Network and the relevant information. The upper panel shows the involvement of miRNAs’ target genes in the enriched pathways (grey girds). The top annotation bars indicate whether the genes are cancer-related, as reported in the two databases COSMIC and OncoKB. Pathways are ranked from top to bottom based on their adjusted p-values in the right annotation, and the left annotation shows the number of involved DEMGIs followed by the number of involved MGIs in the pathways. Genes are ranked from left to right based on ascending log2 fold-change in their expression levels (tumor vs. health), where the stars indicate whether the change is statistically significant (|log2 fold-change| ≥ 1 and adjusted p-value ≤ 0.05). The lower panel in the heatmap shows the top 10 miRNAs with the highest number of identified DEMGIs that are involved in the enriched pathways. The left annotation shows the number of DEMGIs in enriched pathways followed by the number for DE targets in enriched pathways, the number of total DEMGIs, and the number of total target genes. The right annotation shows the results of differential expression analysis including miRNAs’ log2 fold-change and the corresponding adjusted p-values. The grid shows the MGIs’ scores, and the purple frame indicates whether a MGI is a DEMGI. The green annotation bars for miRNAs indicate whether the miRNA are found to be cancer related as reported in the database CMC. **(E)** The Sankey plot illustrates the evidence of miRNA-gene-pathway regulation in BLCA, derived from references (green band) and the Edge-Network results (orange band). We present supporting evidence from literature when published studies examine the same gene predicted by the Edge-Network, and when these studies share either the miRNA or the cancer type with the computational results. Upregulated and downregulated miRNAs and genes in BLCA are colored in pink and blue, respectively. The full data used for the plot can be found in Table S9. LUAD: lung adenocarcinoma; BLCA: bladder urothelial carcinoma; CRC: colorectal cancer.

Furthermore, we search for literature to find evidence for validating the identified 71 DEMGIs and their involvement in regulating cancer phenotypes (Table S8). Among the 33 genes targeted by the top miRNAs, 14 genes (e.g., GNAL, LIFR, PTPRD, SLIT2, MAP1B, and LDLRAD4) are targeted by multiple miRNAs, showing the miRNAs’ redundant and cooperative mechanism for regulating gene expression (9, 10, 64, 65). GNAL functions as a tumor suppressor in glioma by promoting tumor apoptosis, inhibiting tumor proliferation (66), and being associated with clinicopathological features, the tumor immune microenvironment, and therapeutic responses (67). Therefore, the GNAL-targeting miRNAs may function as oncomiRs that promote tumor progression; however, the role of GNAL in BLCA tumorigenesis remains unknown. MiR-182 and miR-93 promote invasion and metastasis in lung adenocarcinoma by repressing LIFR, whereas circCRIM1 counteracts this effect by sponging these miRNAs and restoring LIFR expression (68). The miR-183 cluster, which comprises miR-183, miR-96 and miR-182, is often coordinately expressed and co-regulates multiple pathways in cancers (69). Our results suggest similar regulatory mechanisms in BLCA, where the upregulation of miR-93-5p and miR-183-5p can promote sustained angiogenesis, tissue invasion and metastasis by downregulating LIFR. PTPRD functions as a tumor suppressor in multiplex urothelial carcinomas of the bladder, indicated by its frequent homozygous deletions, biallelic inactivation, recurrent mutations and epigenetic silencing, and the identification of minimal overlapping deleted regions (70), but its regulation by miRNAs remains unknown and we identify six potential miRNAs that can repress its expression in BLCA. SLIT2, which is predicted to be targeted by five miRNAs, promotes tumor growth and invasion in chemically induced skin carcinogenesis (71), but its regulation by miRNAs in BLCA remains unclear. Upregulation of miR-21 downregulates RECK expression, which promotes invasion, proliferation, and migration of high-grade tumor cells in BLCA (72). Our results show that miR-4778-3p and miR-183-5p can also suppress RECK expression, suggesting cooperative miRNA regulation of tumor growth and angiogenesis that promotes tumor progression. MAP1B is overexpressed in bladder cancer and correlates with advanced tumor stage, grade, lymph node metastasis and vascular invasion, while its knockdown reverses chemoresistance by interrupting the cell cycle (73). Our results suggest that MAP1B is involved in regulating tissue invasion and metastasis and can be downregulated by four miRNAs in BLCA. LDLRAD4 functions as both tumor suppressor and oncogene in early hepatocarcinogenesis and hepatic cancer (74, 75). Although LDLRAD4’s role in BLCA remains uncharacterized, our results suggest that four miRNAs can repress its expression, which is associated with the regulation of tumor growth.

We also identify many genes that are targeted by fewer or a single miRNA, suggesting the miRNAs’ specific targeting and regulatory functions in BLCA (Figure 4D). EBF1, a documented cancer driver gene targeted by miR-98-5p, promotes proliferation, migration, and invasion in BLCA through enhanced TMPO-AS1 transcription (76). Our results show that EBF1 could also be downregulated by let-7g-3p, suggesting an alternative regulatory mechanism that may contribute to tumors’ capability to evade growth suppressors. PDE4D is a tumor suppressor in BLCA, with low expression associated with tumor progression and poor prognosis through reduction of intracellular cAMP that enhances the antitumor effect of IFN-α (77). Consistent with this tumor-suppressive role, we identify miR-103a-2-5p as a potential regulator whose upregulation may contribute to PDE4D downregulation in BLCA. SOCS3 has been widely reported as a tumor suppressor and is known to be regulated by various miRNAs across different cancers (78). Specifically, in colon cancer, upregulation of miR-183-5p directly downregulates SOCS3 to upregulate PD-L1 expression, promoting tumor progression by activating the JAK2/STAT3 signaling pathway (79). Our analysis shows that upregulation of miR-183-5p can suppress the tumor suppressor SOCS3, thereby regulating multiple cancer hallmarks in BLCA. DCUN1D3 is a tumor suppressor and is downregulated by miR-93-5p, and its loss promotes tumor proliferation, migration, and invasion in BLCA (80). We show that DCUN1D3 can also be downregulated by miR-183-5p. The oncogene PDCD1LG2, which promotes immune evasion, alters the tumor immune phenotype (81), and enhances metastasis via c-Src/FAK signaling (82), has been shown to be repressed by miR-138-5p in BLCA. Our results suggest that PDCD1LG2 can also be targeted by miR-301b-3p in BLCA, and that this regulation may contribute to tumor’s evasion of growth suppressors, tumor’s tissue invasion and metastasis, and tumor-promoting inflammation.

Based on the analysis above, we identify six target genes (LIFR, EBF1, RECK, SOCS3, DCUN1D3, and PDCD1LG2) that are supported by literature evidence, where the referenced studies investigate the same gene and share either the miRNA or cancer type with our results (Figure 4E). These correspond to eight MGIs from our analysis, which represent promising candidates for experimental validation in BLCA. In addition, we identify novel MGIs that have been minimally studied in BLCA. For instance, miR-4665-5p may downregulate multiple clustered protocadherins (PCDHs), including PCDHGC5, PCDHGB3, PCDHGA12, PCDHGA5, PCDHGA9, PCDHGB7, and PCDHGC3. PCDHs are frequently hypermethylated in multiple solid cancers (83) and exhibit downregulation in BLCA, where they have been associated with tissue invasion and metastasis. The downregulation of PCDHs by miR-4665-5p represents a potential regulatory mechanism that warrants further investigation in BLCA.

Taken together, we demonstrate that the edge-based methods in miREA can identify experimentally validated MGIs supported by the literature, as well as reveal novel MGIs. This provides hypotheses for investigating the molecular mechanisms through which miRNAs regulate gene expression biology and cellular phenotypes in cancer.

## 4 DISCUSSION AND CONCLUSION

We present miREA, a network-based miRNA-oriented enrichment analysis framework. miREA contains five new edge-based algorithms that integrate expression and interactome profiles with pathway networks, providing a more comprehensive representation of miRNA-gene regulatory mechanisms. To our knowledge, this is the first tool that uses network edges for enrichment analysis, with the aim for mitigating the bias of node-based methods that consider only miRNA-gene associations but not the quantitative information in their regulatory interactions. Compared to traditional methods, the edge-based methods show higher sensitivity for curated pathways with biological relevance (i.e., cancer-related pathways), well-controlled errors for randomly shuffled data (i.e., negative pathways), stronger ability to distinguish cancer-specific pathways, remarkable interpretability in identifying context-specific miRNAs and genes, and reasonable computational efficiency after using parallel computing. Furthermore, as demonstrated using BLCA data, miREA can help researchers select suitable methods for their analysis. The results can unravel novel miRNA-mediated gene regulatory program at the pathway level for experimental validation.

Despite its overall satisfactory performance, miREA is subject to several objective limitations that may affect the reliability of its results. First, miREA integrates multiple data sources, including gene expression profiles, pathway annotations, and background MGI and GGI networks. Consequently, its outputs are inherently dependent on the quality, completeness, and credibility of these datasets. Previous studies have shown that ranking metrics, such as Edge-Score in miREA, and the choice of pathway database can significantly impact enrichment results (60, 84). Despite comprehensive curation, the combined MGI and GGI networks in miREA only approximate the true cellular interactome. Condition-specificity, limited experimental validation, and the prevalence of spurious or functionally neutral interactions preclude a complete and precise representation of functional molecular interactions (85). Second, emerging evidence indicates that miRNA-mediated gene regulation is more complex than previously assumed (86, 87). Novel binding mechanisms, including seedless binding (87), cooperative binding (9, 10, 88), and supplementary region modifications (89), as well as the modulatory roles of protein co-factors such as TNRC6C (90, 91) and miRNAs’ AGO-loading status (92), cannot be fully captured by the MGI dataset in miREA. Third, although miRNAs may interact with numerous transcripts at the sequence level, only a subset of these interactions produce measurable biological effects (93). Therefore, although miREA uses experimentally supported and comprehensively curated MGI datasets, it cannot guarantee that all included MGIs are functionally active under the modeled biological conditions. Addressing these limitations would require corresponding data that is not yet available.

The absence of well-established gold-standard benchmarks specifically designed for miRNA-oriented enrichment analysis presents a fundamental challenge for rigorous performance evaluation. Although benchmark frameworks for gene set enrichment analysis have been proposed (59, 94, 95), such as GSEABenchmarkR that assesses method applicability, gene set prioritization, and relevant process detection using microarray and RNA-seq data (95). These frameworks are not directly transferable to miRNA-oriented enrichment methods due to inherent differences in data structure and pathway annotation schemes.

Furthermore, enrichment methods frequently exhibit trade-offs across evaluation metrics, such that no single method uniformly outperforms alternatives across all assessment criteria. To address this issue, we use a thorough benchmarking framework to systematically evaluate edge- and node-based methods in miREA from multiple, complementary perspectives. This approach provides a balanced characterization of the strengths and limitations of each method. This multi-criteria evaluation is intended to help users select the most appropriate method for their data type and analytical objective.

Estimating the false positive rate to assess specificity presents a particular methodological challenge because there is no universally accepted approach for generating biologically meaningless negative controls in the context of miRNA enrichment analysis. Although prior studies have evaluated false positive rates using randomly generated pathways (30, 31, 96), this strategy is not directly applicable here. This is because node-based and edge-based methods operate on different types of input data, such as a list of genes, miRNAs, or MGIs. This makes randomized pathway sets non-comparable across different methods in miREA. To address this issue, we implement a label randomization strategy that shuffles the MGI data in the global MGI list and maps the synthetic MGIs into cancer-specific pathways (53, 59, 97). Compared to the phenotype permutation approach used in most studies, which involves shuffling normal and tumor samples prior to differential expression analysis to generate synthetic MGIs and DEMGIs (94, 95), our approach offers two key advantages. First, it avoids the recomputation of DEMGIs and the introduction of confounding factors. Specifically, the number of DEMGIs derived from permuted phenotypes may differ from those used in the original analysis. This difference can inflate or deflate positive enrichment rates (Figure S9). Second, our approach preserves the integration of the biological network and the empirical distribution of edge scores by retaining the MGI data from the original analysis, thereby avoiding distributional shifts. Furthermore, the proposed methods consider only miRNAs as gene regulators, which simplifies the gene regulatory mechanism. Although transcription factors also play a major role in gene regulation (98), incorporating miRNA regulation by transcription factors into the pathway networks yield negligible effects on the results (data not shown). This is likely attributable to the limited number of experimentally verified transcription factors known to regulate miRNA expression. Together, these properties ensure that the resulting synthetic pathways serve as biologically meaningless negative controls while maintaining analytical consistency with the original enrichment framework at an affordable computational cost.

Although miREA is designed for miRNA-oriented enrichment analysis, we show that integrating gene expression data improves true positive rates (Figure S10). This improvement likely reflects the gene-centric nature of the pathway annotations used in this study. Mapping pathways to their targeting miRNAs introduces an indirect association that may inaccurately represent a miRNA’s function in a pathway. Although miRNA-centric pathway databases, such as miRTar (99), miRPathDB (100), DIANA-miRPath (25), TAM (26) and HMDD (101), may be more appropriate for enrichment analysis using only miRNA expression data, we don’t incorporate them in the benchmarking test for two reasons. First, gene-centric pathway annotations are more widely used and enable direct comparability across enrichment methods, such as node-based methods. Second, these pathways typically derive miRNA-pathway associations through interactions between miRNAs and genes, most of which are predicted by algorithms and not experimentally validated. This can introduce biases that require further empirical evaluation. Nevertheless, using miRNA-centric pathway databases may affect the performance of enrichment analysis. These databases may be better suited for scenarios in which only miRNA data are available; this requires further investigation.

In summary, miREA offers new methods for miRNA-based enrichment analysis and is a valuable addition to the study of cellular miRNA function.

## Authors’ contributions

**ZZ**: methodology; software; data curation; investigation; validation; formal analysis; visualization; project administration; resources; writing – original draft; writing – review and editing. **XL**: conceptualization; methodology; software; data curation; investigation; validation; formal analysis; supervision; funding acquisition; visualization; project administration; resources; writing – original draft; writing – review and editing.

## Data availability

The code and data used in this work are archived in Zenodo (https://doi.org/10.5281/zenodo.18803209). The code and documentation can be found at the GitHub repository https://github.com/laixn/miREA.

## Declaration of AI usage

After writing up the manuscript, we used Claude Sonnet 4.6 to improve the text’s readability and check grammar errors.

## Acknowledgements

We thank the Finnish CSC for providing computational resources and support.

